# Cortical E-fields of deep brain stimulation in Parkinson’s disease patients exceed typical E-field magnitudes of transcranial electrical stimulation

**DOI:** 10.64898/2026.06.06.730577

**Authors:** Thomas Keizers, Axel Thielscher, Oula Puonti, Alessandro Gulberti, Monika Pötter-Nerger, Maria Carla Piastra, Bettina Schwab

## Abstract

**Background:** Deep brain stimulation (DBS) is an established and effective intervention for Parkinson’s disease. Although the exact mechanisms of action are still unclear, both therapeutic benefits and side effects are solely attributed to neuromodulation via strong electric fields (E-fields) in the surgical target. Nevertheless, DBS generates E-fields that extend beyond the stimulation site and can be detected throughout the brain. Recent evidence from transcranial electrical stimulation studies shows that weak cortical E-fields even below 1 V/m can have a neuromodulatory effect, raising the question of a physiological relevance of weak cortical fields of DBS. However, the strength of cortical E-fields of DBS is currently unknown.

**Objective:** Using a novel framework, we aimed to quantify the whole-brain E-field distribution in patients with Parkinson’s disease receiving therapeutic DBS of the subthalamic nucleus (STN-DBS).

**Methods:** In this work, we developed a pipeline to simulate DBS E-field distributions throughout the brain, based on the existing open-source toolboxes Lead-DBS and SimNIBS. We constructed patient-specific whole-head models including electrode leads for 25 patients with Parkinson’s disease receiving subthalamic DBS, and simulated E-fields using the patients’ clinical stimulation settings for 49 hemispheres.

**Results:** We found that median peak E-field magnitudes exceeded 0.3 V/m in all cortical regions and were greater than 1 V/m in the orbital gyrus, superior temporal gyrus, fusiform gyrus, parahippocampal gyrus, insular gyrus, cingulate gyrus. Most prominently, the orbital and insular gyri showed peak magnitudes ranging from 0.81 to 8.61 V/m and 0.90 to 8.01 V/m.

**Conclusions:** Our results indicate that the E-fields of STN-DBS reach cortical peak magnitudes that exceed typically reported values of transcranial electrical stimulation. This opens the possibility that weak E-fields of DBS could have a direct neuromodulatory effect in wider regions of the brain, for example in cortical regions.

## Introduction

Deep Brain Stimulation (DBS) of the subthalamic nucleus (STN) or globus pallidus pars interna (GPi) is an established surgical therapy for Parkinson’s disease (PD). Even though DBS is clinically used in more than 200,000 patients worldwide (1), the exact therapeutic mechanism as well as the mechanism of side effects remain unclear.

Current hypotheses for the mechanism of action pose that purely the stimulation of the target nuclei in the basal ganglia and the white matter bundles surrounding these nuclei are modulated directly. On a cellular level, DBS has been proposed to cause action potentials on axons which travel both ortho- and antidromically. Orthodromic action potentials reach efferent targets and via neurotransmitter release influence postsynaptic potentials. Antidromic action potentials may collide with action potentials originating from the cell body, leading to termination, or reach the cell body thereby affecting the neurons intrinsic firing pattern. Afferent inhibitory axons might also generate orthodromic action potentials, increasing GABAergic signalling leading to increased inhibition. Additionally, activation of nearby white matter fibre bundles and afferent cortical projections could also lead to propagation of action potentials to wide spread regions of the brain, further influencing cortical networks at large (2,3).

Furthermore, DBS has been found to downregulate beta band (13 – 30 Hz) power (4) and beta burst durations (5) within the STN, which have been correlated with reduction of symptom severity. Finally, at the circuit level DBS has been found to alter functional connectivity in PD across motor cortical and subcortical regions exhibiting far reaching network effects (6,7). These hypotheses state that purely the focal stimulation with strong E-fields in or directly around the surgical target drives the therapeutic and side effects. Notably, strong E-fields above 200 V/m can directly activate neurons and axons and therefore form the most prominent stimulation effect within the so-called volume of tissue activated (VTA) (8,9).

In contrast, E-fields outside the VTA are typically neglected. However, from recent studies with transcranial electric stimulation (tES) and more specifically, transcranial alternating current stimulation (tACS) in non-human primates, we know that neuromodulation can be achieved even at much lower field strengths. These weaker, subthreshold E-fields do not, in contrast to suprathreshold E-fields, affect the spike rates of neurons, but rather spike timing preferences of active neurons (10,11). Also in humans, tACS has been shown to have phase-specific neuromodulatory effects, also by influencing connectivity between different cortical regions depending on ongoing oscillatory activity (12–18). Thus, it is possible that these weak E-fields of DBS could also have a neuromodulatory effect that adds on top of the primary effect in the VTA and subsequent spreading throughout the network.

To estimate the relevance of weak E-fields of DBS, the first critical step is to know about their distribution and magnitudes across the whole head. So far, these magnitudes are unknown and difficult to measure directly. For tES, computational whole-head model estimates of E-fields are common and have been validated with intracranial EEG measurements in human epilepsy patients (19). The open-source simulation software package SimNIBS, for example, can be used for such computational studies in non-invasive brain stimulation (20), allowing users to create personalized head models based on magnetic resonance imaging (MRI) and simulate transcranial direct current stimulation to study the distribution of the resulting E-fields over the cortex. For studies on DBS, Lead-DBS (21) is a commonly used tool, which allows users to reconstruct DBS electrode lead trajectories based on pre-operative MRIs and post-operative computed tomography (CT) scans. Users can locate electrode leads and stimulation contacts with respect to subcortical atlases and simulate the VTA to estimate the range of neural activation.

However, to investigate the E-field distribution of DBS across the whole brain, patient-specific whole-head models which also include DBS electrode leads are required. Therefore, we introduce a framework that combines the desired functions of SimNIBS and Lead-DBS to create a head model based on the patients’ individual anatomy and accurately position the DBS electrode leads and contacts. Using finite-element method (FEM) simulations in this framework, we estimated the whole-head E-fields of STN-DBS in a cohort of PD patients and evaluated the distribution of these fields in cortical regions.

## Methods

### Data acquisition and ethical approval

We used previously obtained preoperative T1- and T2 weighted MRIs and post-operative CT-scans for 23 patients (22) in addition to two newly included patients recruited at the University Medical Centre Hamburg-Eppendorf. For each patient, we acquired the clinical stimulation settings and the implanted electrode lead. All 25 patients were diagnosed with PD and received monopolar DBS in the subthalamic nucleus (STN). Ethical permission for the use or re-use and transfer of the defaced, pseudomized data was given by the local ethics committee of the Medical Association of Hamburg (2023-101217-BO-ff).

### Head model creation

An overview of the head model creation process is depicted in Figure 1. We started with the segmentation of the head based on the pre-operative T1- and T2-weighted MRIs using SimNIBS. We then reconstructed the electrode trajectory based the pre-operative T1- and T2-weighted MRIs and post-operative CT-scans using Lead-DBS. Afterwards, we created a voxelized mask of the electrode surface mesh based on the trajectory and a 3D model created in Blender 5.0 (23), and combined this with the upsampled segmentation of the head. This updated segmentation was used to create a whole-head model, which was finalized by including the electrode contacts. All code used in this paper has been made available online (https://gitlab.utwente.nl/bss_development/neuro/dbs-whole-head-modelling/dbs-whole-head-modelling).

**Figure 1:**
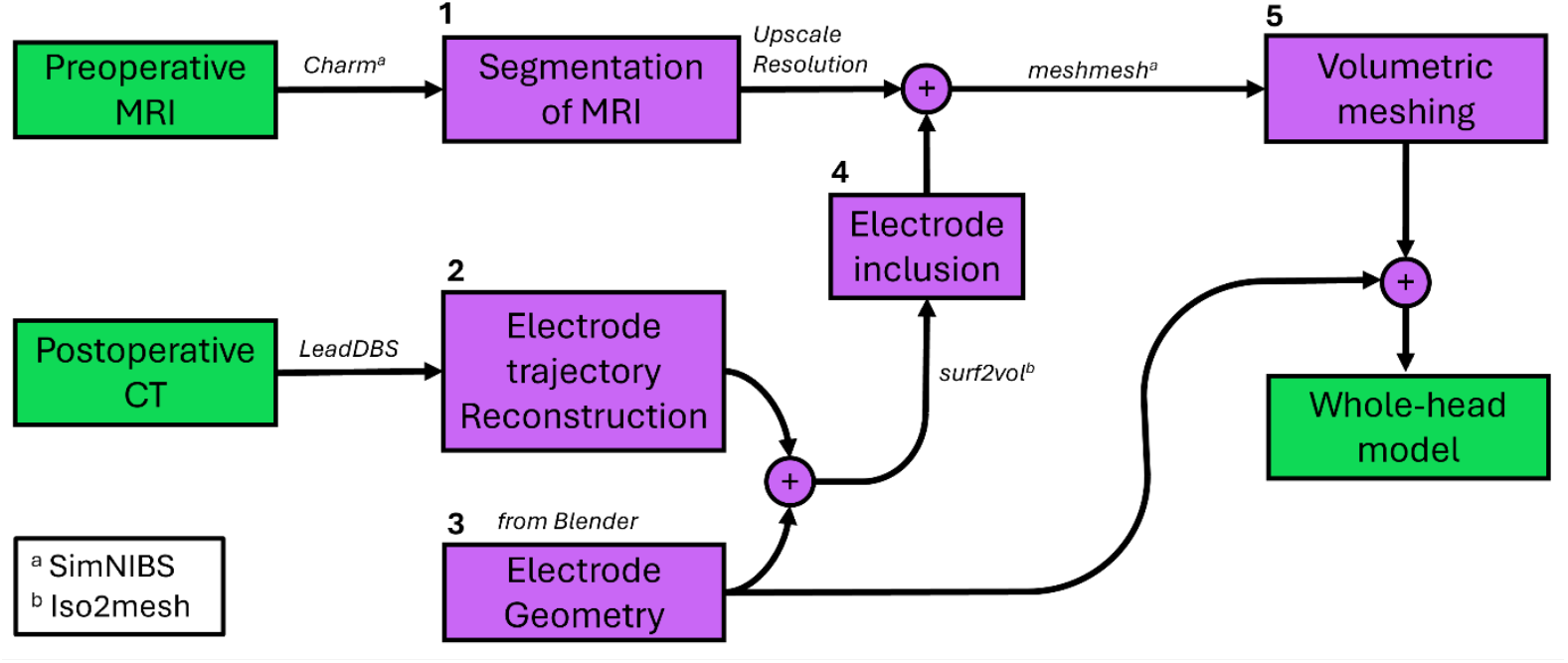
Creation of patient-specific head models from imaging data.

### Segmentation of MRI

We used the *charm* segmentation algorithm in SimNIBS 4.6.0 (20,24) using pre-operative T1- and T2-weighted MRIs to obtain the following tissue compartments: white matter, grey matter, cerebrospinal fluid, scalp, eye balls, compact bone, spongy bone, blood, and muscle.

### Electrode trajectory reconstruction

We obtained the electrode trajectories and contact positions from Lead-DBS (version 3.2.1) (4) based on the patients’ pre-operative T1- and T2-weigthed MRIs and post-operative CT-scans. Images were coregistered, the volumes were normalized, and brainshift correction was applied, the results of which were inspected visually to confirm validity. Following automatic pre-reconstruction the electrodes were localized and the trajectory was obtained in native space.

### Electrode geometry

With Blender (23), we created surface meshes for the following electrode: Medtronic 3389, Medtronic SenSight B33005, Boston Scientific Vercise, and St. Jude Medical 6146_6149 (25–28). The electrode surface meshes were created according to specifications of the electrode geometry on the producer design sheets and constituted the lead shaft and contacts. We modelled the ring electrode contacts as full cylinders, while the segmented contacts were modelled as a hollow cylinder divided into three equivalent segments spanning 100 degrees of the circumference, with 20 degrees spacing between the segments. A schematic drawing can be found in Figure 2, which also shows our adapted contact labelling scheme that will be used throughout this paper for consistency. The number of contacts, contact length, contact spacing and electrode diameter can be found in Table 1.

**Figure 2:**
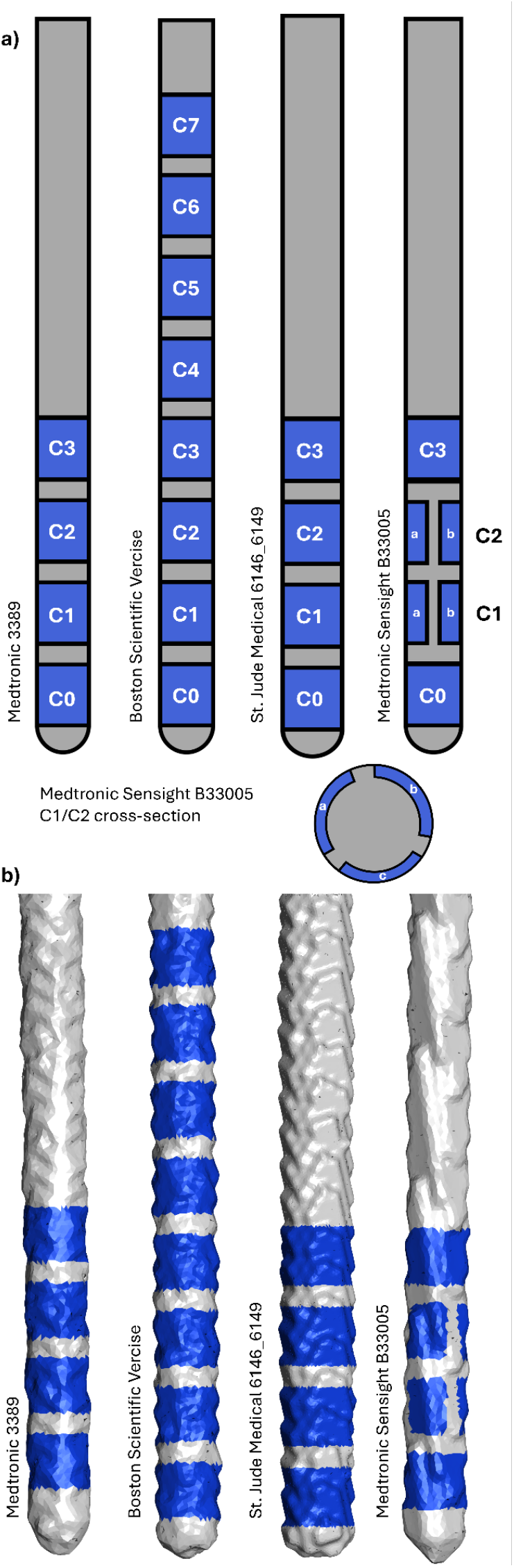
a) Schematic drawings of the electrode models. The insulation and contacts are depicted as grey and blue, respectively. As contact nomenclature differs per electrode model, we defined the most distal contact as C0 for consistency. All models have full ring electrodes, except the Medtronic Sensight B33005, where C1 and C2 have segmented contacts, for which a cross-section with segment labels is provided. b) Volume mesh representations of the electrode models as present in the whole-head models.

**Table 1:** Electrode geometry parameters.

|  | *Medtronic 3389* | *Medtronic Sensight B33005* | *Boston Scientific Vercise* | *St. Jude Medical 6146_6149* |
| --- | --- | --- | --- | --- |
| *Number of contacts* | 4 | 8 | 8 | 4 |
| *Contact length (mm)* | 1.5 | 1.5 | 1.5 | 1.5 |
| *Contact spacing (mm)* | 0.5 | 0.5 | 0.5 | 0.5 |
| *Electrode diameter (mm)* | 1.27 | 1.36 | 1.3 | 1.4 |

### Electrode inclusion

To combine the head segmentation with the electrode geometries, we used the upsampled segmentation output from the charm algorithm. This segmentation consisted of the voxelated label image at a uniform resolution of 0.25 mm to accurately depict the electrode lead with a diameter between 1.27 mm and 1.4 mm. We added an encapsulation layer to allow for different electrophysiological properties at the electrode-tissue interface (29). The encapsulation layer was created by inflating the insulation surface geometry by 0.5 mm in all directions (30). The encapsulation and insulation surfaces were positioned according to the electrode trajectories obtained from Lead-DBS for both hemispheres, voxelated at the resolution of the segmentation and overlaid onto the segmentation image as new labels. The insulation terminated at the boundary between compact bone and scalp, and the encapsulation at the boundary between cerebrospinal fluid and compact bone.

### Volumetric meshing

We created the whole-head model using SimNIBS meshing functionality with the updated segmentation image as input, tissue specific mesh refinement settings can be found in Table S1. The finalized whole-head models contained on average 4,062,806 nodes and 22,587,442 tetrahedra.

The personalized whole-head models included insulation and encapsulation elements, while the contact volume could not be included in the segmentation images due to resolution constraints with respect to the contact spacing. Therefore, the contact elements were included in the volumetric model by relabelling insulation elements. This was done by aligning the electrode surface geometries with the insulation volume and relabelling those elements which geometric centres fell within the closed surface of the electrode contacts. The mesh refinement within the insulation shaft was high enough to ensure the contacts were separated by multiple tetrahedra. The shared boundaries between the newly labelled contacts and the encapsulation served as input locations for the stimulation.

For monopolar stimulation, a return electrode outside of the electrode lead is required. In the human body, the implanted pulse generator (IPG) serves this purpose, implanted below the clavicle, which falls outside of the domain of our head models. Therefore, we approximated the location by creating a surface patch at the base of the neck 3 cm to the right of the spinal column, yielding a consistent IPG approximation location between individuals. This surface patch served as the output location for the currents injected on the electrode contacts.

### Mesh quality metrics

We investigated the quality of our head models by calculating three element quality metrics, namely, the Signed Inverse Condition Number (SICN), the ratio of the inscribed radius and the circumscribed radius (gamma), and the signed inverse error on the gradient of FE solution (SIGE), as provided by gmsh (31). All three metrics ranged from 0, i.e. completely degenerate elements, to 1, i.e. ideally proportioned elements. Furthermore, the number of elements of low quality, defined as elements with quality metrics <0.1, was quantified. To check the effect of including the electrode geometries on the overall mesh quality, we compared these metrics between the head mesh provided by the initial run of the charm algorithm and the head mesh created from the upsampled new segmentation file obtained.

### Patient information

We included data of 25 patients with monopolar stimulation from other clinical studies at the Neurology Department of the University Medical Center Hamburg-Eppendorf. Inclusion criteria were: Diagnosis of Parkinson’s disease, monopolar DBS of the subthalamic nucleus, availability of preoperative T1- and T2-weighted MRIs without imaging artefacts, postoperative CT-scans, stimulation settings, and electrode model. The electrode model, active contacts, input amplitudes, and input type for each patient are summarized in Table 2. This information was used to construct the patient-specific head models and to define their given input parameters for the simulation.

**Table 2:** Patient stimulation settings.

| Patient | Electrode model | Input type | Right hemisphere |  | Left hemisphere |  |
| --- | --- | --- | --- | --- | --- | --- |
|  |  |  | Contacts | Input ampl. | Contacts | Input ampl. |
| Pat1 | Vercise | mA | C3 | 3.8 | C3 | 3.4 |
| Pat2 | Med3389 | V | C0 / C1 | 2.9 / 2.9 | C0 / C1 | 3.6 / 3.6 |
| Pat3 | Med3389 | V | C1 | 3.2 | C1 | 3.2 |
| Pat4 | Med3389 | V | C1 / C2 | 2.6 / 2.6 | C1 / C2 | 2 / 2 |
| Pat5 | Med3389 | V | C2 | 2 | C2 | 3.8 |
| Pat6 | Vercise | mA | C2 / C3 | 1.5 / 1.5 | C2 / C3 | 1.75 / 1.75 |
| Pat7 | Med3389 | V | C2 | 2.7 | C0 / C1 | 2.5 / 2.5 |
| Pat8 | Med3389 | V | C1 / C2 | 2.5 / 2.5 | C1 / C2 | 2.5 / 2.5 |
| Pat9 | Vercise | mA | C4 | 4.5 | C3 | 4 |
| Pat10 | Vercise | mA | C3 | 3.4 | C3 | 3 |
| Pat11 | Med3389 | mA | C2 | 3.5 | C1 | 1.7 |
| Pat12 | Med3389 | V | C1 / C2 | 2.3 / 2.3 | C1 / C2 | 1.2 / 1.2 |
| Pat13 | Med3389 | mA | C3 | 1.5 | C3 | 2 |
| Pat14 | Vercise | mA | C2 | 2.1 | C2 | 2.2 |
| Pat15 | Vercise | mA | C2 / C3 | 1.5 / 1.5 | C3 | 3.6 |
| Pat16 | Abbott | mA | C2 / C3 | 1 / 1 | C1 / C2 | 1.85 / 1.85 |
| Pat17 | Med3389 | V | - | - | C1 | 1.6 |
| Pat18 | Med3389 | V | C1 / C2 | 3.6 / 3.6 | C1 / C2 | 2 / 2 |
| Pat19 | Vercise | mA | C1 / C3 | 1.55 / 1.55 | C1 / C3 | 1.45 / 1.45 |
| Pat20 | Abbott | mA | C1 / C2 | 1.25 / 1.25 | C1 / C2 | 1.25 / 1.25 |
| Pat21 | Med3389 | mA | C2 | 3.3 | C2 | 2.7 |
| Pat22 | Vercise | mA | C2 | 3.8 | C2 | 3.5 |
| Pat23 | Med3389 | V | C1 | 2.9 | C1 | 2.9 |
| Pat24 | Vercise | mA | C2 | 2.6 | C1 | 2.3 |
| Pat25 | B33005 | mA | C2 b/c | 1.2 / 1.2 | C2 a/b/c | 0.77 / 0.77 / 0.77 |

### Simulations

To calculate the resulting E-field distribution, we used the FEM functions provided by SimNIBS. The tissue conductivities were based on the standard conductivities used in SimNIBS for the common tissue types and encapsulation (29,32–34) are provided in Table 3. The conductivities for the insulation and contacts were based on the material properties of polyurethane (35) and platinum-iridium (36) respectively. The conductivity of the insulation was increased from 10^−14^ S/m to 10^−6^ S/m, as even lower values led to problems with convergence.

**Table 3:** Conductivities of tissues and materials.

| Material/tissue type | Conductivity (S/m) |
| --- | --- |
| White Matter | 0.126 |
| Grey Matter | 0.275 |
| Cerebrospinal Fluid | 1.654 |
| Skin | 0.467 |
| Eyeballs | 0.5 |
| Compact Bone | $8 \times 10^{-3}$ |
| Spongy Bone | $2.5 \times 10^{-2}$ |
| Blood | 0.6 |
| Muscle | 0.16 |
| Encapsulation | $1.9 \times 10^{-2}$ |
| Insulation | $1 \times 10^{-6}$ |
| Contact | $4 \times 10^3$ |

Our dataset consists of patients with either current or voltage controlled stimulation. For voltage controlled stimulation, we calculated the current flow at the neck patch to allow for comparison between the two stimulation configurations.

The electric potential field, Φ, was simulated and the E-field was derived as *E* = −∇ϕ. The E-field was calculated at the centre of each tetrahedron.

### Quantification of E-fields

#### Volume of Tissue Activated

To calculate the Volume of Tissue Activated (VTA), we considered the E-field values in the white matter, grey matter, and encapsulation layers. The absolute magnitude of the E-field values was thresholded at 200 V/m and the element volumes summed to obtain the VTA per hemisphere. We also calculated the VTA for all patients using Lead-DBS, via Lead Group’s (32) *Calculate VTA & Stats*, as an additional way to check the quality of our VTA estimations. We used a Wilcoxon signed rank test to test for differences in the estimated VTA.

#### Cortical segmentation

To estimate E-field values in cortical regions of the brain, we aligned the Brainnetome atlas (38) with our head models. We used the closed surface meshes for each atlas region to assign the elements of the head mesh to their corresponding atlas region.

#### Outcome metrics

For E-field vectors within a cortical region, we calculated the means and 99.9^th^ percentiles of the magnitudes, corrected for element size. The 99.9^th^ percentile of all data points was chosen rather than the absolute maximum to avoid excessively high values that could potentially be caused by simulation noise or computational outliers. To calculate the E-field normal component with respect to the cortical surface, we determined which face of the cortical surface was closest to each element and projected the E-field vector onto the normal component. For simplicity, we report all metrics only for the stimulated hemisphere.

For visualization purposes, we transformed the simulation results to the FreeSurfer average template space. This step provided E-fields in the grey matter layer projected onto the inflated cortical surface and allowed for comparison across patients as well as group level visualization.

#### Insular cortex

Finally, we focused on the E-field magnitude distribution in a single cortical region. We chose the insular cortex, which due to its position deep in the lateral sulcus experiences the highest average DBS E-fields compared to other cortical regions.

## Results

### Head models

We obtained patient-specific head models including DBS leads, an example of which can be seen in Figure 3. Then, we calculated mesh quality metrics for each patient-specific head model before and after our meshing processing (supplemental materials S1), indicating a lower number of low-quality elements after processing.

**Figure 3:**
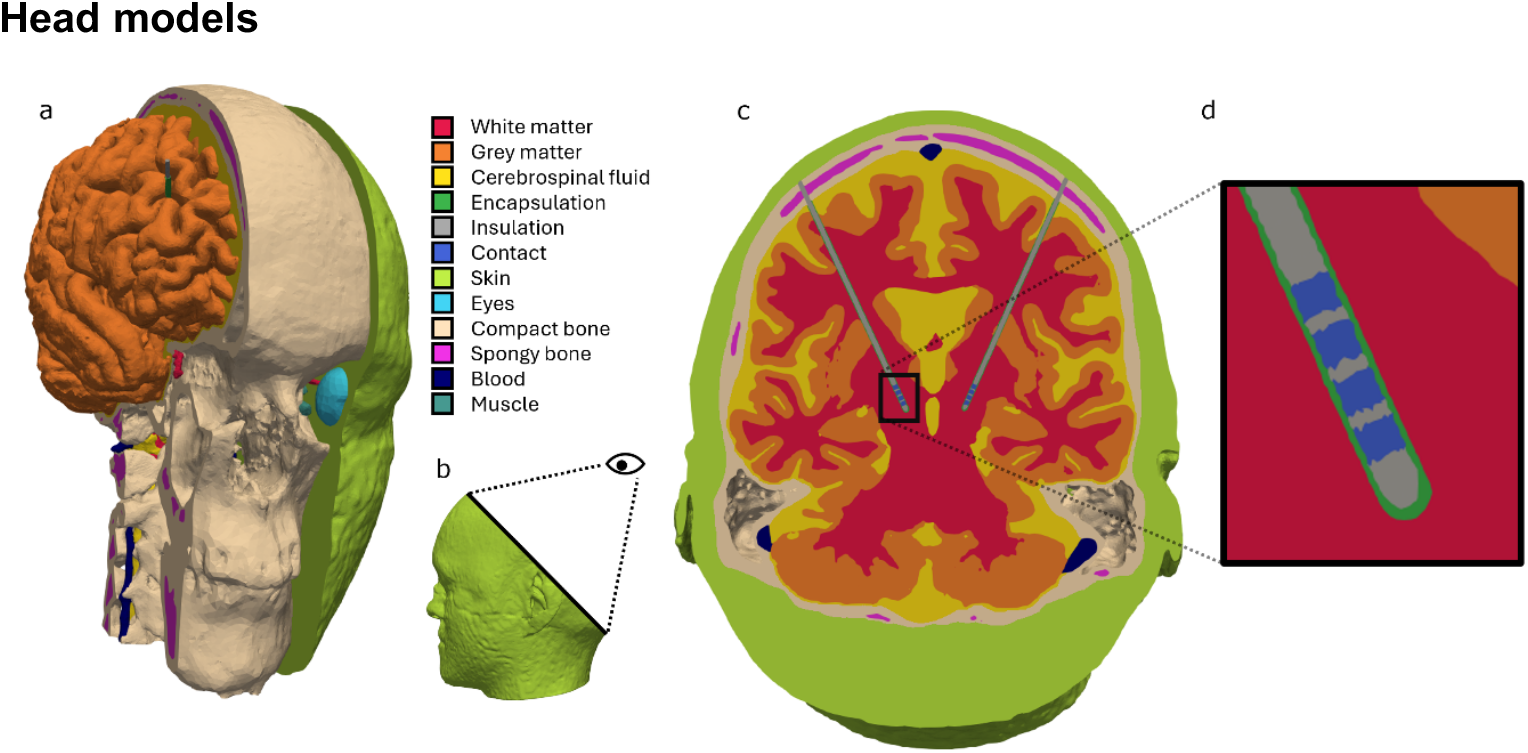
Volume layers of an example head model. a) The head model with volume layers cut at sagittal planes at different points along the frontal axis, showing the brain, skull, and scalp. b) A view of the head model in the sagittal plane showing the orientation of the cutplane and viewpoint. c) The cut plane through the head model along the axis of the electrode leads. d) The electrode lead and encapsulation layer, zoomed in on the same cutplane.

### Simulations

#### Voltage-controlled stimulation

The voltage input and the corresponding current input per hemisphere are shown in Table 4 for all patients who received voltage-controlled stimulation.

**Table 4:** The calculated input current for patients receiving voltage-controlled stimulation. All patients receiving voltage-controlled stimulation were implanted with Medtronic 3389 electrodes.

| Patient | Hemisphere | Contact(s) | V | mA |
| --- | --- | --- | --- | --- |
| Pat2 | Right | C0 + C1 | 2.9 | 2.70 |
|  | Left | C0 + C1 | 3.6 | 4.32 |
| Pat3 | Right | C1 | 3.2 | 1.73 |
|  | Left | C1 | 3.2 | 3.82 |
| Pat4 | Right | C1 + C2 | 2.6 | 3.08 |
|  | Left | C1 + C2 | 2.0 | 1.79 |
| Pat5 | Right | C2 | 2.0 | 1.83 |
|  | Left | C2 | 3.8 | 3.44 |
| Pat7 | Right | C2 | 2.7 | 1.51 |
|  | Left | C0 + C1 | 2.5 | 2.26 |
| Pat8 | Right | C1 + C2 | 2.5 | 3.53 |
|  | Left | C1 + C2 | 2.5 | 2.94 |
| Pat12 | Right | C1 + C2 | 2.3 | 2.69 |
|  | Left | C1 + C2 | 1.2 | 1.41 |
| Pat17 | Left | C1 | 1.6 | 1.93 |
| Pat18 | Right | C1 + C2 | 3.6 | 3.41 |
|  | Left | C1 + C2 | 2.0 | 1.80 |
| Pat23 | Right | C1 | 2.9 | 1.58 |
|  | Left | C1 | 2.9 | 1.66 |

### Quantification of E-fields

#### Volumes of tissue activated are comparable to Lead-DBS

Lead-DBS reconstructions (Figures 4a and 4b) showed that the electrodes of all patients successfully reached the target site. The active contacts were all within or near the boundary of the STN.

**Figure 4:**
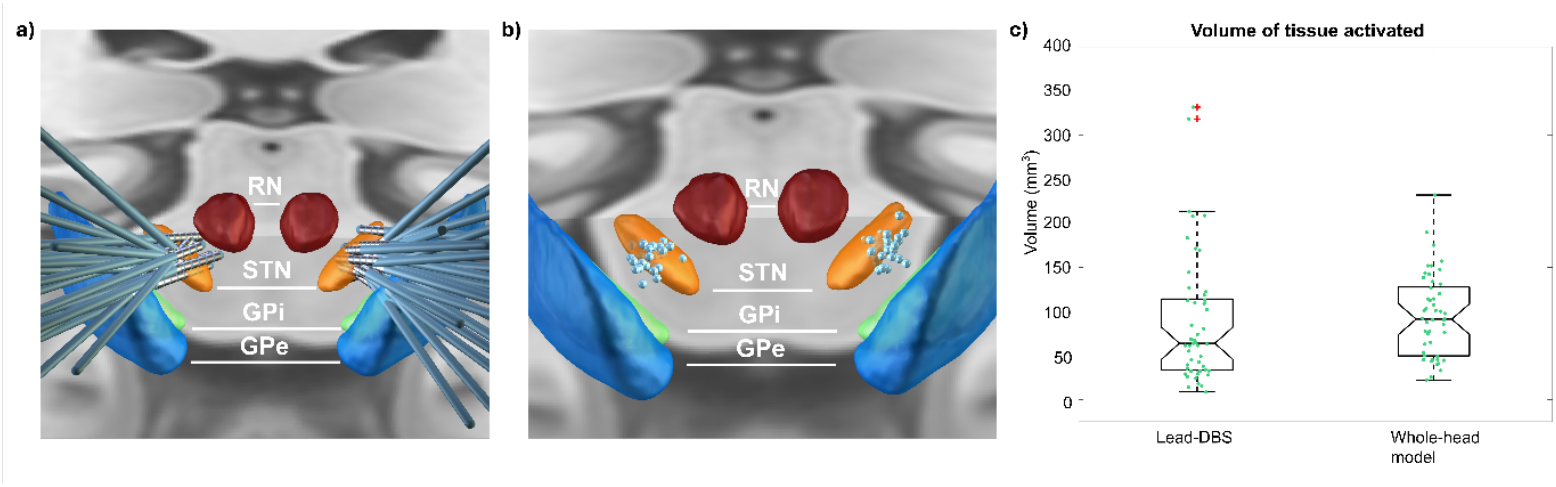
a) Group visualization of the electrode trajectories in Lead-DBS. (RN = red nucleus, STN = subthalamic nucleus, GPi = globus pallidus pars interna, GPe = globus pallidus pars externa) b) Group visualization of the active contacts. The points cluster around the STN, the stimulation target. c) The VTA estimates for both Lead-DBS and our SimNIBS-based whole-head model. The VTA estimates for all patients for both hemispheres (n=49) were pooled together and shown as points along the boxplots. The 25^th^ and 75^th^ quartiles are indicated by the box, the whisker are given by 1.5 times their respective interquartile lengths. The notch around the median indicates the 95% confidence interval of the estimated median. Outliers are points that fall outside of the whiskers and are marked with red crosses.

To investigate the E-fields in the immediate surroundings of the electrode, we calculated the VTA for all hemispheres with both Lead-DBS and our whole-head model (Figure 4c). The mean volume as calculated by Lead-DBS was 87.88 +/- 73.75 mm^3^ and the mean volume obtained from the whole-head model was 94.74 +/- 46.58 mm^3^. We observed no significant group-level difference between the VTAs obtained from Lead-DBS and the whole head model according to the Wilcoxon signed rank test (p = 0.303).

#### E-fields in the cortex exceed 1 V/m

The E-field magnitude of the group average at the inflated cortical surface can be seen in Figure 5a, for both hemispheres from lateral and medial viewpoints per stimulated hemisphere. As expected, the magnitudes were higher in deeper regions of the brain. This can be seen in the depths of the central and lateral sulci. The lower magnitudes at the most superior parts of the cortex were due to the larger distance to the stimulating contacts compared to the more lateral parts, which can be seen in Figure 5b.

**Figure 5:**
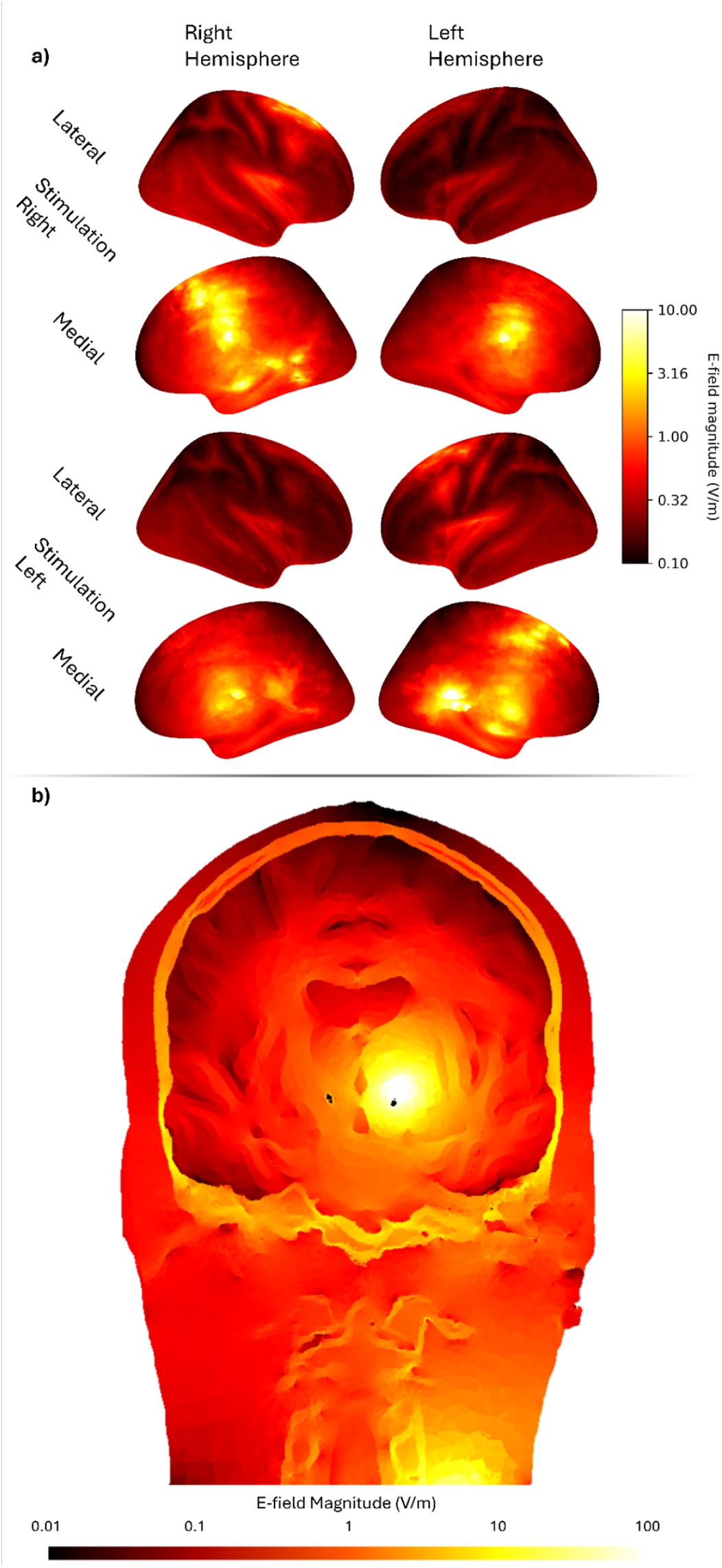
a) Group average in the FreeSurfer average template space projected onto the inflated cortical surface for stimulation in the left and right hemispheres. b) Cross-section of the head model for Pat 1, cut along the coronal plane viewed posteriorly. The total stimulation amplitude was 3.8 mA applied to contact C3 in the right hemisphere.

We summarized the mean and 99.9^th^ percentile magnitudes of the E-field in the Brainnetome atlas regions of each hemisphere (Figures 6a and 6b, respectively). The lowest mean value was found in the superior parietal lobule, with a median value across hemispheres of 0.10 V/m (Figure 6a). The highest mean values were found in the parahippocampal and insular gyri, with medians across hemispheres of 0.89 V/m and 0.95 V/m, respectively.

**Figure 6:**
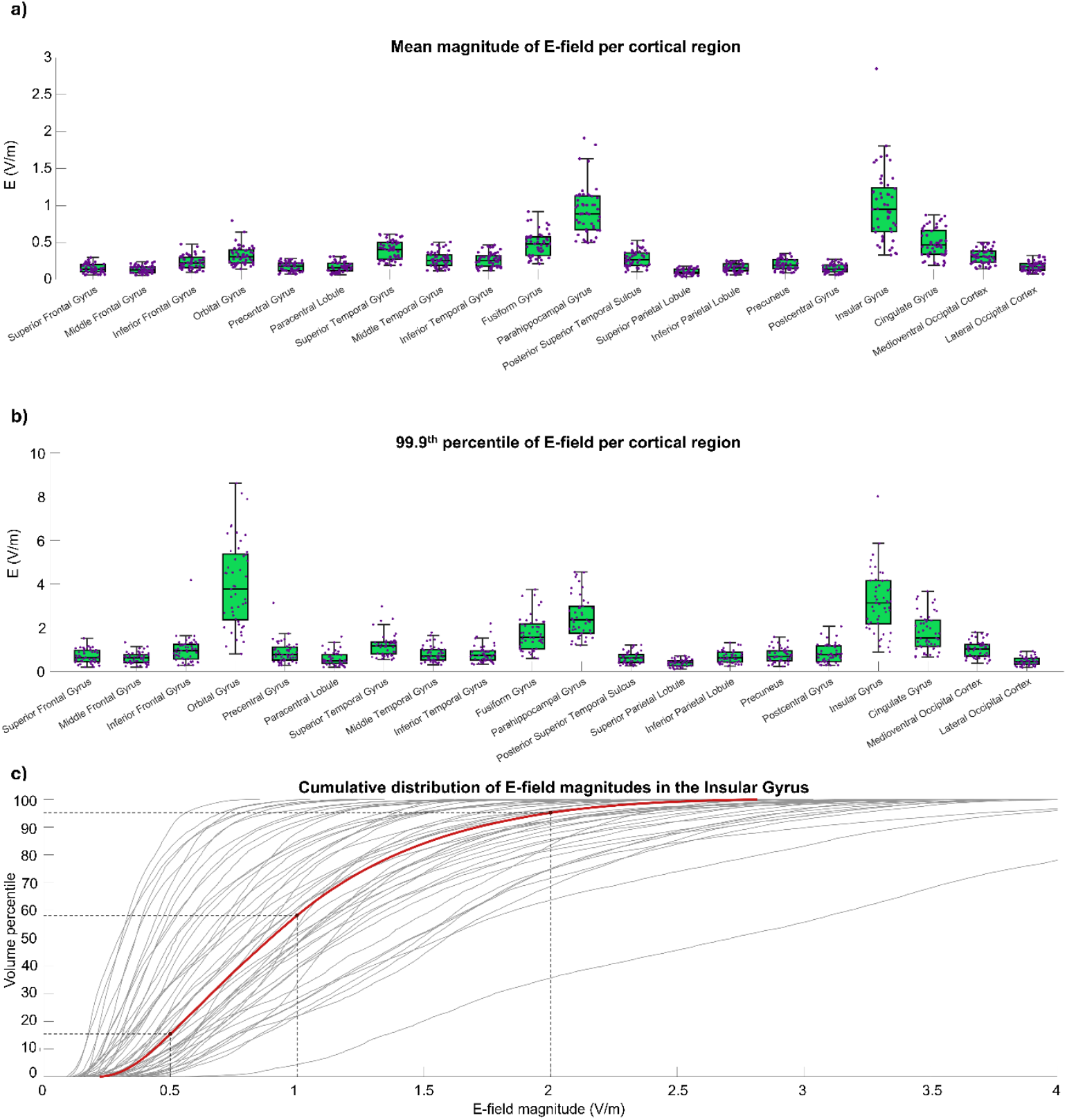
E-field magnitudes reached in 20 cortical regions for all hemispheres. The median value across hemispheres is indicated by the red line, the box represents the 25^th^ and 75^th^ percentiles of the distribution. An outlier is indicated by a red plus mark and is at least 1.5 times the interquartile distance above or below the boundaries of the box. A) Mean values of the E-field magnitude reached in each cortical region. B) 99.9^th^ percentile values of the E-field magnitude reached in each cortical region. C) E-field magnitudes reached in the insular cortex of the stimulated hemisphere for all 49 simulated hemispheres. The individual hemispheres are depicted in grey and the average E-field magnitude in red.

Figure 6b shows the 99.9^th^ percentile “peak” value of the E-field for each patient in the specified cortical region. Its median values across hemispheres exceeded 0.5 V/m in all regions, except the paracentral lobule, superior parietal lobule, and lateral occipital cortex. Six regions had median 99.9^th^ percentile values exceeding 1 V/m, being the orbital gyrus, superior temporal gyrus, fusiform gyrus, parahippocampal gyrus, insular gyrus, cingulate gyrus. In general, the highest values were reached in the orbital and insular gyri reaching medians of 3.77 V/m and 3.16 V/m, respectively. Furthermore, the peak E-field magnitudes in the orbital gyrus exceeded 5 V/m in 15/49 hemispheres and the insular gyrus reached peak E-field magnitudes above 5 V/m in 4/49 hemispheres.

The E-field magnitudes achieved in the insular gyrus showed strong differences between hemispheres/patients (Figure 6c). For example, the magnitudes achieved in the most stimulated 0.01% volume ranged from 0.85 to 6.76 V/m. The lowest values reached in the insula gyrus ranged from 0.09 V/m to 0.56 V/m. On average, the lowest magnitude in the insular gyrus was 0.22 V/m up to a 99.9^th^ percentile of 2.82 V/m. On average, the values of 0.5 V/m, 1.0 V/m, and 2.0 V/m were reached at the 15.5^th^, 58.2^th^, and 95.2^th^ percentiles.

## Discussion

In this study, we combined features of existing open source toolboxes to create accurate whole-head models of patients with DBS implants. Using this novel framework, we were able to simulate the E-fields of DBS throughout the whole-head model for a cohort of 25 PD patients. We quantified E-fields in the local, subcortical environment and also throughout cortical regions, demonstrating that cortical E-fields of DBS reach strengths that are comparable to or higher than typical E-field strengths of tACS. As tACS can have considerable phase-specific effects on both neurophysiology and behaviour (39–41), our results highlight the possibility of direct neuromodulation by DBS far outside of the VTA, including cortical regions.

While this study is the first to quantify cortical E-fields of DBS, it is not the first to use larger volume conduction models to simulate E-fields of DBS. For example, OSS-DBS increased the simulation space from a limited domain around the electrode to a whole-brain domain (42). Nevertheless, there are key differences between our goals and methods. We aimed to gain information on E-field distribution in distinct cortical regions for patient models using clinical stimulation settings. In contrast, OSS-DBS is more focused on simulating in the local electrode environment to quantify the direct activation of neurons surrounding the active contacts. Compared to OSS-DBS, our head model consists of the entire head using personalized anatomy, rather than brain tissues surrounded by cerebrospinal fluid. Furthermore, our head models can also be used in simulations of other brain stimulation techniques, such as transcranial electrical stimulation, which would allow us to investigate distributions during multimodal stimulation configurations like simultaneous DBS and tACS.

Another tool developed for FEM simulations of electrical stimulation in larger head models is FEMfuns (43). FEMfuns is able to perform simulations for resistive, capacitive and dispersive regimes, and can also take anisotropy into account. Both OSS-DBS and FEMfuns are capable of simulating E-fields of DBS in larger head models and include expanded simulation capabilities compared to what we show here. We chose to perform the simulations in SimNIBS because its existing segmentation pipeline enabled an accurate whole-head model creation process. We used SimNIBS segmentation and meshing procedures which are already tuned for creating models to simulate in SimNIBS, exporting and adjusting volumetric models for different simulation environments adds further complexity. Furthermore, performing simulations within SimNIBS provides easy access to additional post-processing and visualization functionalities. Thus, as we did not intend to investigate capacitive or dispersive effects, we opted for SimNIBS as the final FEM solver for a more streamlined process.

Our framework is generic and can be used for any DBS indication, for example also essential tremor or epilepsy. The only requirement is the availability of preoperative MRIs for the segmentation and postoperative CT or MRI for electrode trajectory reconstruction. All code used for head model creation in this paper has been made publicly available. To ensure the code can be tested, we provide sample images and data based on the publicly available Ernie dataset (20).

As a first validation of the E-field estimates for DBS, we calculated the VTA with our approach and compared it with the VTA computed in Lead-DBS. The VTA as used here is a simple, heuristic outcome metric, which in recent years is used more often in combination with tractography and connectomics (8). The methods of calculation are straightforward, and the comparison demonstrated that the E-field estimation in the immediate surroundings of the electrode was comparable to Lead-DBS, strengthening the validity of our model.

Having validated the E-field estimations in the local electrode environment, we proceeded to investigate the distribution of cortical DBS E-fields. The magnitude of the cortical E-fields was comparable to field strengths reported in human tACS literature or higher. Specifically, tACS typically leads to peak E-field strengths below 1 V/m peak amplitude (14,17,18,44–49). In general, the field strengths present in the cortex during DBS within our patient population exceeded the theoretical minimum effective dose of 0.3 V/m as proposed by Alekseichuk et al. (50) for tACS, with the lowest median peak magnitude being 0.40 V/m, in the superior parietal lobule. Single neuron measurements in macaques have shown entrainment effects of tACS at starting at maximum E-field magnitudes of 0.3 – 0.5 V/m (10,11). We find that cortical E-fields of DBS are at least as large and may even be ten times larger, depending on the region. For example, median peak magnitudes exceeding 3 V/m were found in the orbital and insular gyri.

We specifically focused on the insular cortex as it is an important site for sensory integration, and plays a large role in affective, cognitive and sensorimotor networks. The insular cortex is also one of the most strongly stimulated cortical regions in our model, experiencing peak magnitudes from 0.90 up to 8.01 V/m. The posterior insular cortex has connections to premotor areas of the cortex and electrical stimulation via intracortical microstimulation has been found to generate hand movements in macaques (51). In humans, the posterior insular cortex functionally connects to motor cortical areas (52). Furthermore, the insular cortex shows increased functional magnetic resonance imaging (fMRI) activity during STN-DBS (53). Additionally, a functional connection between the STN and the insular cortex was also found in both healthy adults and PD patients using arterial spin-labelled perfusion fMRI (53).

Together, these findings stress a potential physiological role of the insular cortex and its connectivity to the basal ganglia for DBS.

Nevertheless, it is still unclear how the response of cortical neurons to the E-fields of DBS and tACS compare. Even though the magnitudes of the E-fields in cortical areas for DBS compared to tACS might be higher, e.g. in the insular and cingulate cortices, there are also differences between frequencies and waveshapes. For tACS, the stimulation frequency is dependent on the frequency band of interest but commonly spans from theta (4 – 8 Hz) up to gamma (30 – 100 Hz) frequencies (54–56). The stimulation frequency for STN-DBS in PD is typically in the range of 60 - 130 Hz, with 130 Hz being the most often used frequency (57). Low frequency stimulation (<100 Hz) can be used to treat axial symptoms more specifically (58). Furthermore, tACS often uses sinusoidal waves, but DBS waveshapes consist of short pulses. Given the nonlinear nature of neuronal networks, the response to different waveforms can strongly differ. Thus, modeling and experimental studies that investigate the influence of E-fields in the determined range of a few V/m with waveshapes of short pulses are required to understand the functional consequences of the cortical E-fields of DBS.

Additionally, most DBS devices do not apply stimulation for both hemispheres simultaneously, but the stimulation is provided in an interleaved fashion. This is particularly relevant in subcortical regions roughly equidistant from both electrodes. Depending on the orientation of the neurons relative to the E-fields originating from both active contacts, some subcortical regions could experience stimulation at twice the stimulation frequency. This is the case for the entire brain, as the interleaving of pulses means effectively stimulating at twice the frequency, but depending on the exact regions the differences in field strengths between the stimulation originating from the two hemispheres might cause one source to be too weak to have an effect. In sum, depending on the distance to the stimulation sites and the orientation compared to the generated E-fields, some neurons could be experiencing E-fields above 1 V/m at double the DBS frequency. The functional consequences of such stimulation still need to be explored.

In this study, we found that STN-DBS E-fields on average reach maximum values above 0.5 V/m in the majority of cortical regions. This indicates large portions of the brain are perpetually exposed to E-fields exceeding the minimum effective dose of tACS, with deeper regions, such as the insular and cingulate cortices, experiencing values that can even be one order of magnitude greater. There is a possibility that DBS could therefore affect cortical regions directly via weak E-fields rather than purely via direct and indirect synaptic connections with the basal ganglia. In particular, if entrainment occurs sporadically in cortical areas due to weak E-fields and frequently in subcortical areas due to strong E-fields, certain functional cortico-subcortical connections may specifically be enhanced or depressed. Critically, cortical E-fields showed a strong inter-patient variability, which could potentially explain part of the inter-participant variability in clinical DBS outcomes or side effects. Further studies are needed to investigate the functional and clinical consequences of these weak cortical E-fields of DBS.

### Limitations

Our model is based on previously established and validated methods, the core features and parameter settings were not altered. Nevertheless, the application of SimNIBS to DBS is new and not directly experimentally validated. A possible addition to the model would be anisotropic tissue properties based on the tractography from diffusion tensor imaging, which SimNIBS provides. Also, we did not account for spatial variation in electrical properties within the compartments in addition to anisotropy (58). Still, our model accurately reproduces VTAs as computed with Lead-DBS, supporting its overall validity.

Furthermore, inclusion of the encapsulation layer was based on findings of changes in the tissue surrounding the electrode (28,29,59–61), with electrical properties estimated from previously reported values. However, there is variation in the conductivity values reported for different tissue types and we assumed these values to be constant within their associated tissues (62). In the future, it may be possible to estimate personalized tissue conductivities by measuring the DBS artefacts on scalp EEGs and obtaining the values via optimization procedures.

Finally, in this study, we focused on DBS of the STN for Parkinson’s disease. Different E-field distributions may be observed for other DBS targets and indications, requiring different stimulation amplitudes. Importantly, stimulation amplitudes for Parkinson’s disease are not at the higher end of the spectrum. Indications like dystonia (64), epilepsy (65), obsessive-compulsive disorder (66), and depression (67) often require higher amplitudes, which will also lead to higher E-field magnitudes in cortical areas as these scale linearly with stimulation amplitude. Therefore, our estimates for Parkinson’s disease represent rather a lower bound of E-field values that are expected for DBS in general, although the exact distributions across cortical sites may differ.

## Conclusion

In conclusion, using our new framework, we were able to simulate the E-field distribution throughout the whole brain during DBS for a cohort of PD patients. On average, maximum E-field magnitudes exceed 1 V/m in several cortical regions. Deeper cortical regions such as the insular and parahippocampal gyri may experience field strengths of at least 2.5 V/m depending on the individual stimulation settings and patient anatomy. Taken together, E-field magnitudes in the cortex during DBS exceed reported magnitudes typically reached by tES.

## CRediT authorship contribution statement

Thomas Keizers: Methodology, Software, Validation, Formal Analysis, Investigation, Data Curation, Writing – Original Draft, Visualization

Axel Thielscher: Methodology, Writing – Review and Editing

Oula Puonti: Methodology, Writing – Review and Editing

Alesandro Gulberti: Resources, Writing – Review and Editing

Monika Pötter-Nerger: Resources, Writing – Review and Editing

Maria Carla Piastra: Methodology, Supervision, Writing – Review and Editing

Bettina C. Schwab: Conceptualization, Resources, Data Curation, Methodology, Supervision, Writing – Review and Editing, Project Administration, Funding Acquisition

## Data and code availability

Data and code for this study are available via Gitlab (https://gitlab.utwente.nl/bss_development/neuro/dbs-whole-head-modelling/dbs-whole-head-modelling)

## Acknowledgements

This work was financed by the European Research Council (ERC StG DECODE, grant number 101116047, to B.C.S). B.C.S. further received funding from the German Research Foundation (DFG, grant number SCHW 2023/2–1) and the Dutch Research Council (NWO, grant number 22332). A.T. was supported by the German Research Foundation (Research Unit 5429/1 (467143400), TH 1330/6-1, TH 1330/7-1) and by the Lundbeck foundation (grant R313-2019-622). O.P. received funding from Lundbeckfonden (grant number R360–2021– 395). M.C.P. received funding from the Dutch Research Foundation (NWO, VENI project 20194). We would like to thank Roger Dozen for his help with management of MRI and CT data, and Johannes Vorwerk for valuable discussions.

## Supplements

### S1. Mesh Element Volumes

**Table S1:** Tetrahedron element size range.

| Tissue label | Minimum size | Maximum size |
| --- | --- | --- |
| White matter | 1 | 5 |
| Grey matter | 1 | 7 |
| Scalp | 1 | 10 |
| Encapsulation | 0.25 | 0.5 |
| Insulation | 0.05 | 0.1 |
| Default | 1 | 5 |

### S2. Mesh Quality Metrics

The average values (mean +/- std) of SICN, gamma, and SIGE for the meshes before any additional processing, i.e. the immediate result of the SimNIBS charm procedure, were 0.7624 +/- 0.0007, 0.7132 +/- 0.0008, and 0.8050 +/- 0.0004 respectively. The average values of SICN, gamma, and SIGE of the meshes resulting from the electrode inclusion pipeline were 0.7465 +/- 0.0007, 0.6981 +/- 0.0009, and 0.8002 +/- 0.0004 respectively. The Wilcoxon signed rank test showed that the electrode inclusion procedure has a statistically significant effect on all of the quality metrics (p < 0.001), with differences (before – after) of 0.0159 (p < 0.001, CI95% = [0.0155, 0.0163]) for SICN, 0.0151 (p < 0.001, CI95% = [0.0147, 0.0155]) for gamma, and 0.0047 (p < 0.001, CI95% = [0.0046, 0.0049]) for SIGE.

The number of low-quality elements (quality metric < 0.1) for SICN, gamma, and SIGE were 707 +/- 112, 3176 +/- 447, and 681 +/- 109 respectively before any additional processing. After electrode inclusion the amount of low-quality elements for SICN, gamma, and SIGE were 726 +/- 98, 4,615 +/- 991, and 689 +/- 87 respectively. However, the total number of elements differed between the two stages increasing from 4,155,340 pre-processing to 22,587,442 post-processing. Accordingly, fraction of low-quality elements decreased from 1.70 * 10^-4 to 3.21 * 10^-5 for SICN, from 7.64 * 10^-4 to 2.04 * 10^-4 for gamma, and from 1.64 * 10^-4 to 3.05 * 10^-5 for SIGE.

To summarize, while there was a very small deterioration of average mesh quality with our new procedure, the amount of low-quality elements relative to the total amount of elements was decreased.

### S3. Normal components of cortical E-fields

**Figure S1:**
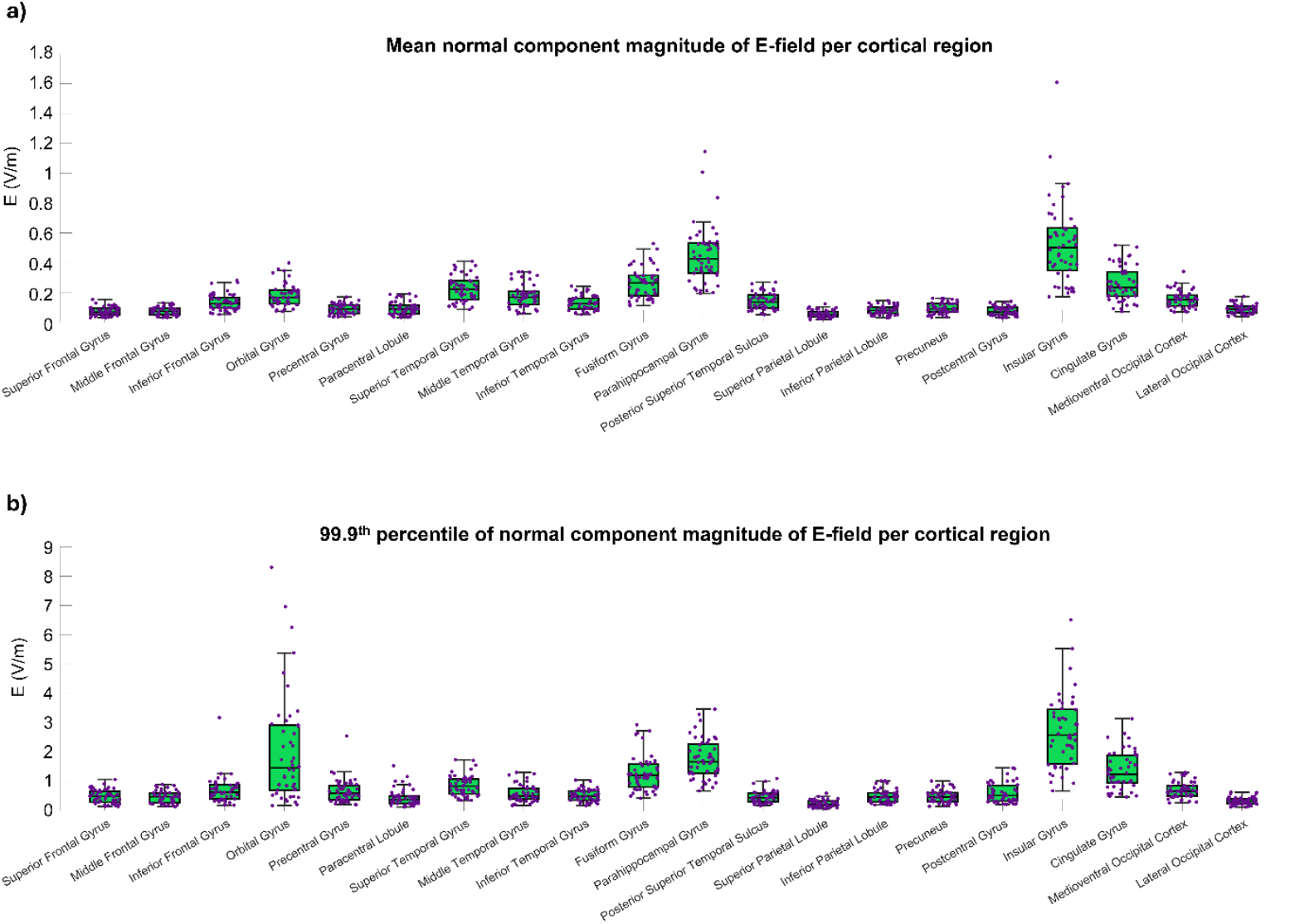
E-field normal component magnitudes with respect to the cortical surface reached in 20 cortical regions for all hemispheres. The median value across hemispheres is indicated by the red line, the box represents the 25^th^ and 75^th^ percentiles of the distribution. An outlier is indicated by a red plus mark and is at least 1.5 times the interquartile distance above or below the boundaries of the box. A) Mean values of the E-field normal component magnitude reached in each cortical region. B) 99.9^th^ percentile values of the E-field normal component magnitude reached in each cortical region.

## References

1. Vedam-Mai V, Deisseroth K, Giordano J, Lazaro-Munoz G, Chiong W, Suthana N, et al. Proceedings of the Eighth Annual Deep Brain Stimulation Think Tank: Advances in Optogenetics, Ethical Issues Affecting DBS Research, Neuromodulatory Approaches for Depression, Adaptive Neurostimulation, and Emerging DBS Technologies. Front Hum Neurosci. 2021 Apr 19;15. doi:10.3389/fnhum.2021.644593

2. Jakobs M, Fomenko A, Lozano AM, Kiening KL. Cellular, molecular, and clinical mechanisms of action of deep brain stimulation—a systematic review on established indications and outlook on future developments. EMBO Mol Med. 2019;11(4). Located at: Scopus. doi:10.15252/emmm.201809575

3. Neumann WJ, Steiner LA, Milosevic L. Neurophysiological mechanisms of deep brain stimulation across spatiotemporal resolutions. Brain. 2023 Nov 1;146(11):4456–68. doi:10.1093/brain/awad239

4. Binns TS, Köhler RM, Vanhoecke J, Chikermane M, Gerster M, Merk T, et al. Shared pathway-specific network mechanisms of dopamine and deep brain stimulation for the treatment of Parkinson’s disease. Nat Commun. 2025 Apr 15;16(1):3587. doi:10.1038/s41467-025-58825-z

5. Anidi C, O’Day JJ, Anderson RW, Afzal MF, Syrkin-Nikolau J, Velisar A, et al. Neuromodulation targets pathological not physiological beta bursts during gait in Parkinson’s disease. Neurobiol Dis. 2018 Dec 1;120:107–17. doi:10.1016/j.nbd.2018.09.004

6. Bove F, Genovese D, Moro E. Developments in the mechanistic understanding and clinical application of deep brain stimulation for Parkinson’s disease. Expert Rev Neurother. 2022;22(9):789–803. Located at: Scopus. doi:10.1080/14737175.2022.2136030

7. Horn A, Wenzel G, Irmen F, Huebl J, Li N, Neumann WJ, et al. Deep brain stimulation induced normalization of the human functional connectome in Parkinson’s disease. Brain. 2019;142(10):3129–43. Located at: Scopus. doi:10.1093/brain/awz239

8. Patrick EE, Fleeting CR, Patel DR, Casauay JT, Patel A, Shepherd H, et al. Modeling the volume of tissue activated in deep brain stimulation and its clinical influence: a review. Front Hum Neurosci. 2024;18. Located at: Scopus. doi:10.3389/fnhum.2024.1333183

9. Åström M, Diczfalusy E, Martens H, Wårdell K. Relationship between Neural Activation and Electric Field Distribution during Deep Brain Stimulation. IEEE Trans Biomed Eng. 2015 Feb;62(2):664–72. doi:10.1109/TBME.2014.2363494

10. Johnson L, Alekseichuk I, Krieg J, Doyle A, Yu Y, Vitek J, et al. Dose-dependent effects of transcranial alternating current stimulation on spike timing in awake nonhuman primates. Sci Adv. 2020 Sep 2;6(36):eaaz2747. doi:10.1126/sciadv.aaz2747

11. Krause MR, Vieira PG, Csorba BA, Pilly PK, Pack CC. Transcranial alternating current stimulation entrains single-neuron activity in the primate brain. Proc Natl Acad Sci. 2019 Mar 19;116(12):5747–55. doi:10.1073/pnas.1815958116

12. Kim Y, Lee JH, Park JC, Kwon J, Kim H, Seo J, et al. Neuromodulation of inhibitory control using phase-lagged transcranial alternating current stimulation. J NeuroEngineering Rehabil. 2024 May 30;21(1):93. doi:10.1186/s12984-024-01385-y

13. Violante IR, Li LM, Carmichael DW, Lorenz R, Leech R, Hampshire A, et al. Externally induced frontoparietal synchronization modulates network dynamics and enhances working memory performance. Hamilton R, editor. eLife. 2017 Mar 14;6:e22001. doi:10.7554/eLife.22001

14. Schwab BC, Misselhorn J, Engel AK. Modulation of large-scale cortical coupling by transcranial alternating current stimulation. Brain Stimulat. 2019 Sep 1;12(5):1187–96. doi:10.1016/j.brs.2019.04.013

15. Schwab BC, König P, Engel AK. Spike-timing-dependent plasticity can account for connectivity aftereffects of dual-site transcranial alternating current stimulation. NeuroImage. 2021 Aug 15;237:118179. doi:10.1016/j.neuroimage.2021.118179

16. Gann MA, Paparella I, Zich C, Grigoras IF, Huertas-Penen S, Rieger SW, et al. Dual-site beta transcranial alternating current stimulation during a bimanual coordination task modulates functional connectivity between motor areas. Brain Stimulat. 2025 Sep 1;18(5):1566–78. doi:10.1016/j.brs.2025.08.011

17. Fiene M, Schwab BC, Misselhorn J, Herrmann CS, Schneider TR, Engel AK. Phase-specific manipulation of rhythmic brain activity by transcranial alternating current stimulation. Brain Stimulat. 2020 Sep 1;13(5):1254–62. doi:10.1016/j.brs.2020.06.008

18. Fiene M, Radecke JO, Misselhorn J, Sengelmann M, Herrmann CS, Schneider TR, et al. tACS phase-specifically biases brightness perception of flickering light. Brain Stimulat. 2022 Jan 1;15(1):244–53. doi:10.1016/j.brs.2022.01.001

19. Huang Y, Liu AA, Lafon B, Friedman D, Dayan M, Wang X, et al. Measurements and models of electric fields in the in vivo human brain during transcranial electric stimulation. Ivry R, editor. eLife. 2017 Feb 7;6:e18834. doi:10.7554/eLife.18834

20. Thielscher A, Antunes A, Saturnino GB. Field modeling for transcranial magnetic stimulation: A useful tool to understand the physiological effects of TMS? In: 2015 37th Annual International Conference of the IEEE Engineering in Medicine and Biology Society (EMBC) [Internet]. 2015 [cited 2025 Feb 6]. p. 222–5. Available from: https://ieeexplore.ieee.org/document/7318340 doi:10.1109/EMBC.2015.7318340

21. Horn A, Kühn AA. Lead-DBS: A toolbox for deep brain stimulation electrode localizations and visualizations. NeuroImage. 2015 Feb 15;107:127–35. doi:10.1016/j.neuroimage.2014.12.002

22. Schott FP, Gulberti A, Pinnschmidt HO, Gerloff C, Moll CKE, Schaper M, et al. Subthalamic Deep Brain Stimulation Lead Asymmetry Impacts the Parkinsonian Gait Disorder. Front Hum Neurosci. 2022 Mar 28;16. doi:10.3389/fnhum.2022.788200

23. Blender [Internet]. Stichting Blender Foundation. Available from: http://www.blender.org

24. Puonti O, Van Leemput K, Saturnino GB, Siebner HR, Madsen KH, Thielscher A. Accurate and robust whole-head segmentation from magnetic resonance images for individualized head modeling. NeuroImage. 2020 Oct 1;219:117044. doi:10.1016/j.neuroimage.2020.117044

25. Medtronic Activa Tremor Control System [Internet]. Medtronic Inc.; 2002. Available from: https://www.accessdata.fda.gov/scripts/cdrh/cfdocs/cfpma/pma.cfm?id=P960009

26. Vercise Deep Brain Stimulation System Premarket Approval [Internet]. Boston Scientific Corporation; 2017. Available from: https://www.accessdata.fda.gov/scripts/cdrh/cfdocs/cfpma/pma.cfm?ID=P150031

27. Brio Neurostimulation System [Internet]. St. Jude Medical; 2015. Available from: https://www.accessdata.fda.gov/scripts/cdrh/cfdocs/cfpma/pma.cfm?id=p140009

28. Medtronic SenSight Directional Lead Kit Implantation Manual. Medtronic Inc.; 2021.

29. Evers J, Sridhar K, Liegey J, Brady J, Jahns H, Lowery M. Stimulation-induced changes at the electrode–tissue interface and their influence on deep brain stimulation. J Neural Eng. 2022 Jul;19(4):046004. doi:10.1088/1741-2552/ac7ad6

30. Butson CR, Maks CB, McIntyre CC. Sources and effects of electrode impedance during deep brain stimulation. Clin Neurophysiol. 2006 Feb 1;117(2):447–54. doi:10.1016/j.clinph.2005.10.007

31. Gmsh: A 3-D finite element mesh generator with built-in pre- and post-processing facilities -Geuzaine -2009 -International Journal for Numerical Methods in Engineering -Wiley Online Library [Internet]. [cited 2026 Jan 9]. Available from: https://onlinelibrary-wiley-com.ezproxy2.utwente.nl/doi/abs/10.1002/nme.2579?sid=worldcat.org

32. Opitz A, Paulus W, Will S, Antunes A, Thielscher A. Determinants of the electric field during transcranial direct current stimulation. NeuroImage. 2015 Apr 1;109:140–50. doi:10.1016/j.neuroimage.2015.01.033

33. Wagner TA, Zahn M, Grodzinsky AJ, Pascual-Leone A. Three-dimensional head model Simulation of transcranial magnetic stimulation. IEEE Trans Biomed Eng. 2004 Sep;51(9):1586–98. doi:10.1109/TBME.2004.827925

34. Gabriel C, Peyman A, Grant EH. Electrical conductivity of tissue at frequencies below 1 MHz. Phys Med Biol. 2009 Jul;54(16):4863. doi:10.1088/0031-9155/54/16/002

35. Electrical polyurethane properties [Internet]. [cited 2026 Mar 12]. Available from: https://gallaghercorp.com/wp-content/uploads/2016/08/WP-116-ELECTRICAL-POLYURETHANE-PROPERTIES.1.pdf

36. Cardarelli F. Materials Handbook. 2nd edn. Springer London. 1339 p. doi:10.1007/978-1-84628-669-8

37. Treu S, Strange B, Oxenford S, Neumann WJ, Kühn A, Li N, et al. Deep brain stimulation: Imaging on a group level. NeuroImage. 2020 Oct 1;219:117018. doi:10.1016/j.neuroimage.2020.117018 PubMed PMID: 32505698.

38. Fan L, Li H, Zhuo J, Zhang Y, Wang J, Chen L, et al. The Human Brainnetome Atlas: A New Brain Atlas Based on Connectional Architecture. Cereb Cortex. 2016 Aug 1;26(8):3508–26. doi:10.1093/cercor/bhw157

39. Burke R, Maÿe A, Misselhorn J, Fiene M, Engelhardt FJ, Schneider TR, et al. The role of delta phase for temporal predictions investigated with bilateral parietal tACS. Brain Stimulat. 2025 Jan 1;18(1):103–13. doi:10.1016/j.brs.2024.12.1476

40. Fiene M, Schwab BC, Misselhorn J, Herrmann CS, Schneider TR, Engel AK. Phase-specific manipulation of rhythmic brain activity by transcranial alternating current stimulation. Brain Stimulat. 2020 Sep 1;13(5):1254–62. doi:10.1016/j.brs.2020.06.008

41. Fiene M, Radecke JO, Misselhorn J, Sengelmann M, Herrmann CS, Schneider TR, et al. tACS phase-specifically biases brightness perception of flickering light. Brain Stimulat. 2022 Jan 1;15(1):244–53. doi:10.1016/j.brs.2022.01.001

42. Butenko K, Bahls C, Schröder M, Köhling R, van Rienen U. OSS-DBS: Open-source simulation platform for deep brain stimulation with a comprehensive automated modeling. PLoS Comput Biol. 2020 Jul;16(7):e1008023. doi:10.1371/journal.pcbi.1008023 PubMed PMID: 32628719; PubMed Central PMCID: PMC7384674.

43. Vermaas M, Piastra MC, Oostendorp TF, Ramsey NF, Tiesinga PHE. FEMfuns: A Volume Conduction Modeling Pipeline that Includes Resistive, Capacitive or Dispersive Tissue and Electrodes. Neuroinformatics. 2020 Oct;18(4):569–80. doi:10.1007/s12021-020-09458-8 PubMed PMID: 32306231; PubMed Central PMCID: PMC7498500.

44. Leunissen I, Van Steenkiste M, Heise KF, Monteiro TS, Dunovan K, Mantini D, et al. Effects of beta-band and gamma-band rhythmic stimulation on motor inhibition. iScience. 2022 May;25(5):104338. doi:10.1016/j.isci.2022.104338

45. Reinhart RMG. Disruption and rescue of interareal theta phase coupling and adaptive behavior. Proc Natl Acad Sci. 2017 Oct 24;114(43):11542–7. doi:10.1073/pnas.1710257114

46. Zhu T, Sack AT, Leunissen I. Phase-Specific Dual-Site Beta Transcranial Alternating Current Stimulation Differentially Influences Functional Connectivity Associated With Motor Inhibition Performance. Hum Brain Mapp. 2026;47(3):e70470. doi:10.1002/hbm.70470

47. Johari K, Tabari F. Left Frontal HD-tACS Enhances Prefrontal Theta Activity During Action Verbal Fluency in Parkinson’s Disease. Eur J Neurosci. 2026;63(1):e70368. doi:10.1111/ejn.70368

48. Van Hoornweder S, Mora DAB, Nuyts M, Cuypers K, Verstraelen S, Meesen R. The causal role of beta band desynchronization: Individualized high-definition transcranial alternating current stimulation improves bimanual motor control. NeuroImage. 2025 May 15;312:121222. doi:10.1016/j.neuroimage.2025.121222

49. Elyamany O, Iffland J, Bak J, Classen C, Nolte G, Schneider TR, et al. Predictive role of endogenous phase lags between target brain regions in dual-site transcranial alternating current stimulation. Brain Stimulat. 2025 May 1;18(3):780–93. doi:10.1016/j.brs.2025.04.011

50. Alekseichuk I, Wischnewski M, Opitz A. A minimum effective dose for (transcranial) alternating current stimulation. Brain Stimul Basic Transl Clin Res Neuromodulation. 2022 Sep 1;15(5):1221–2. doi:10.1016/j.brs.2022.08.018 PubMed PMID: 36044976.

51. Simone L, Caruana F, Borra E, Del Sorbo S, Jezzini A, Rozzi S, et al. Anatomo-functional organization of insular networks: From sensory integration to behavioral control. Prog Neurobiol. 2025 Apr;247:102748. doi:10.1016/j.pneurobio.2025.102748

52. Cauda F, D’Agata F, Sacco K, Duca S, Geminiani G, Vercelli A. Functional connectivity of the insula in the resting brain. NeuroImage. 2011 Mar 1;55(1):8–23. doi:10.1016/j.neuroimage.2010.11.049

53. Fernández-Seara MA, Mengual E, Vidorreta M, Castellanos G, Irigoyen J, Erro E, et al. Resting state functional connectivity of the subthalamic nucleus in Parkinson’s disease assessed using arterial spin-labeled perfusion fMRI. Hum Brain Mapp. 2015;36(5):1937–50. doi:10.1002/hbm.22747

54. Lee TL, Lee H, Kang N. A meta-analysis showing improved cognitive performance in healthy young adults with transcranial alternating current stimulation. Npj Sci Learn. 2023 Jan 3;8(1):1. doi:10.1038/s41539-022-00152-9

55. Tang Y, Liu D, Li S, Zhu L, Zhu Y, Shi N, et al. Effects of transcranial alternating current stimulation on cognitive function in older adults: a systematic review and meta-analysis of randomized controlled trials. Neurol Sci. 2025;46(12):6227–41. doi:10.1007/s10072-025-08519-7

56. Schutter DJLG, Wischnewski M. A meta-analytic study of exogenous oscillatory electric potentials in neuroenhancement. Neuropsychologia. 2016 Jun 1;86:110–8. doi:10.1016/j.neuropsychologia.2016.04.011

57. Subthalamic Nucleus Deep Brain Stimulation in Parkinson’s Disease: The Effect of Varying Stimulation Parameters -Viswas Dayal, Patricia Limousin, Thomas Foltynie, 2017 [Internet]. [cited 2026 Jan 30]. Available from: https://journals.sagepub.com/doi/10.3233/JPD-171077

58. Mann GS, Panjiyar BK, Allam MP, Asifa F, Ullah A, Jha S. Comparing the impact of low frequency vs high frequency deep brain stimulation in Patient’s with Parkinson’s disease: A systematic review. Brain Disord. 2025 Sep 1;19:100239. doi:10.1016/j.dscb.2025.100239

59. Kumar N, Ahamparam A, Lu CW, Malaga KA, Patil PG. Modeling electrical impedance in brain tissue with diffusion tensor imaging for functional neurosurgery applications. J Neural Eng. 2024 Oct;21(5):056036. doi:10.1088/1741-2552/ad7db2

60. Payonk JP, Bathel H, Arbeiter N, Kober M, Fauser M, Storch A, et al. Improving computational models of deep brain stimulation through experimental calibration. J Neurosci Methods. 2025;414. doi:10.1016/j.jneumeth.2024.110320

61. Kathrin B, Marco S, Thomas K, Eilhard M, Jan G. Impedance detection of the electrical resistivity of the wound tissue around deep brain stimulation electrodes permits registration of the encapsulation process in a rat model. J Electr Bioimpedance. 2017 Apr 1;8(1):11–24. doi:10.5617/jeb.4086

62. Cui J, Mivalt F, Sladky V, Kim J, Richner TJ, Lundstrom BN, et al. Acute to long-term characteristics of impedance recordings during neurostimulation in humans. J Neural Eng. 2024 Apr;21(2):026022. doi:10.1088/1741-2552/ad3416

63. McCann H, Pisano G, Beltrachini L. Variation in Reported Human Head Tissue Electrical Conductivity Values. Brain Topogr. 2019 Sep 1;32(5):825–58. doi:10.1007/s10548-019-00710-2

64. Magown P, Andrade RA, Soroceanu A, Kiss ZHT. Deep brain stimulation parameters for dystonia: A systematic review. Parkinsonism Relat Disord. 2018 Sep 1;54:9–16. doi:10.1016/j.parkreldis.2018.04.017

65. Li MCH, Cook MJ. Deep brain stimulation for drug-resistant epilepsy. Epilepsia. 2018;59(2):273–90. doi:10.1111/epi.13964

66. van Westen M, Rietveld E, Bergfeld IO, de Koning P, Vullink N, Ooms P, et al. Optimizing Deep Brain Stimulation Parameters in Obsessive–Compulsive Disorder. Neuromodulation Technol Neural Interface. 2021 Feb 1;24(2):307–15. doi:10.1111/ner.13243

67. Ramasubbu R, Lang S, Kiss ZHT. Dosing of Electrical Parameters in Deep Brain Stimulation (DBS) for Intractable Depression: A Review of Clinical Studies. Front Psychiatry. 2018 Jul 11;9. doi:10.3389/fpsyt.2018.00302

